# AnimalTA: A simple yet flexible tool for video tracking and manual corrections

**DOI:** 10.64898/2026.06.27.733780

**Authors:** V. Chiara, A. Buatois, S.-Y. Kim

## Abstract

1. Video-tracking programs have now become an essential tool for researchers measuring animal behavior across biological fields. The panel of available programs is growing rapidly, providing researchers with numerous specific tools that will match their precise needs. However, their proliferation may complicate post-tracking data processing, and some programs do not even provide tools for correcting tracking errors or analysing tracking data. In the case of commercial software, the loss of access to a program due to budget limitations or researchers’ mobility from one institution to another could prevent them from accessing and visualizing their tracking data.
2. There is therefore a growing need for an accessible and flexible tool to handle post-tracking processes such as the correction and analysis of tracking data obtained across different video-tracking programs.
3. We present here the latest update of the video tracking and analysis program AnimalTA. With this new release, we propose to solve the above-mentioned problems by providing the scientific community with a program that will allow for data importation from other video-tracking programs. Like in its previous versions, AnimalTA remains a free, open-source, and highly user-friendly program, ensuring that it will always be accessible without restriction. Now, with this new importation option, users who performed their tracking with other programs can benefit from AnimalTA’s complete toolset of data visualization, correction, and analysis.
4. Finally, this article gives an overview of the other main improvements associated with this new release. The program is now faster in both video importation and tracking, proposes an amplified toolset for data visualization and correction, and features new options for data analysis.

## Introduction

The quantitative study of animal behaviour is becoming increasingly reliant on automated tracking technologies that can extract high-resolution movement data from large, complex datasets (Dell et al., 2014). Over the past decade, video-based tracking has become a key methodology in behavioural ecology, ethology, and applied animal science. It enables researchers to capture behavioural phenotypes with a level of precision and efficiency that manual observation cannot match. This methodological shift has been accompanied by a growing demand for transparency, reproducibility, and long-term data accessibility — values that are now central to open science frameworks in the biological sciences (Bell et al., 2009; Wilson et al., 2023). In this context, software tools used for collecting, processing, and analysing behavioural data are foundational components of scientific infrastructure, not merely technical conveniences.

In recent years, the landscape of animal tracking software has diversified considerably, reflecting the range of research questions and experimental systems investigated by behavioural scientists. General-purpose tracking tools, such as AnimalTA, ToxTrack and idTracker, offer robust solutions for tracking multiple individuals under controlled conditions (Chiara & Kim, 2023; Rodriguez et al., 2018; Pérez-Escudero et al., 2014; Torrents et al., 2026). Meanwhile, machine-learning-based pose estimation platforms, such as DeepLabCut (Nath et al., 2019) and SLEAP (Pereira et al., 2022), have transformed our ability to track fine-scale movements of body parts across a wide range of species and contexts. More specialised tools, such as Loopy (http://loopb.io, Loopbio Gmbh, Vienna, Austria), address specific experimental designs or taxa. Despite this diversity, a persistent challenge remains: the output formats, data structures, and downstream analytical workflows associated with these tools are rarely interoperable, limiting the adaptive and flexible use of tracking, correction, and analysis functions of different programs within a given research. Although some programs provide integrated trajectory correction modules, to our knowledge, no tools currently allow manual correction of trajectories independently of their original tracking pipelines, making it difficult to obtain fully verified, functional datasets. Researchers working across platforms or seeking to correct and reanalyse existing datasets often encounter significant barriers to data integration and reuse.

Open-source, freely available software plays a critical role in overcoming these barriers, particularly for researchers working outside well-resourced institutional environments. A recurring practical challenge in this area concerns early-career researchers and students who conduct data collection within a host laboratory, but then lose access to proprietary or institutionally licensed tracking software when they change position or affiliation. This means that previously obtained raw tracking data cannot be curated, corrected, or reanalysed without incurring the cost of accessing the original software environment, posing a significant obstacle to research continuity and reproducibility. Free, open-access tools with broad compatibility therefore serve not only scientific purposes but also promote equity within the scientific community, ensuring that analytical capacity is not contingent on institutional affiliation or grant availability.

AnimalTA was developed in response to these needs, providing a free, user-friendly, open-source platform for tracking and analysing the movement and behaviour of individual or grouped animals (Chiara & Kim, 2023). AnimalTA can track the movement of one or multiple individuals across one or multiple arenas, visualize and correct the resulting trajectories, and analyze these trajectories to extract parameters such as total distance traveled, average speed, path meander, explored area, time spent in a selected area, and average inter-individual distance over time and across time sequences. Since its initial publication, the software has been widely adopted within the animal science community, with over 7,140 downloads and 56 citations reflecting a genuine demand for accessible, well-performing analytical tools. The original version of AnimalTA was primarily designed to work with data generated by its own tracking engine. However, user feedback and evolving community practices have highlighted the need to extend its compatibility to data produced by other widely used tracking platforms and enhance the interactive data curation and error correction tools available.

Here, we present a substantial update to AnimalTA that directly addresses these needs. The updated software can now import and process tracking outputs from both general-purpose and machine-learning-based tracking tools, such as ToxTrack, DeepLabCut, SLEAP, and Loopy. It has also been designed to be extensible, so that it can accommodate additional formats as the needs of the community evolve. New interactive functionalities have been implemented for tracking data correction and curation, providing real-time visual feedback that facilitates the rapid identification and repair of tracking errors. These features are particularly valuable in teaching and training contexts. These developments position AnimalTA as a powerful, shared analytical environment that bridges tracking platforms, reduces pipeline fragmentation, and supports the reproducibility and accessibility of behavioural research, regardless of the upstream tools used.

## Presentation of the new features of AnimalTA

### 1. General presentation

AnimalTA is an open-source, Python-developed video-tracking program released for the first time in 2023 (Chiara & Kim, 2023). It uses various libraries, including tkinter for the Graphical User Interface (GUI), OpenCV for image processing, ffmpeg, and decord for video conversion and reading. AnimalTA uses a user-friendly Windows OS installer built using Inno Setup (https://jrsoftware.org/isinfo.php) and can be downloaded along with a user manual and tutorial videos at: http://vchiara.eu/index.php/animalta.

The previous version of AnimalTA was already a fully functional program, whose characteristics have been described in Chiara & Kim (2023). Below are the main new features and improvements of the program to import, curate, correct, and analyse animal tracking data originating from other tools.

### 2. Loading trajectory data from other video-tracking programs

With this update, AnimalTA now supports researchers who obtain tracking data using tools other than AnimalTA but seek a faster and more efficient way to visualize, correct, and analyze their data. It is also useful for those who no longer have access to the initial tools, which were accessible thanks to an institutional licence. The updated program includes a new “Import Data” panel, which allows users to combine previously imported videos with individual and group trajectory data from external sources.

Prior to importing trajectory data, users must first upload their videos into the program using the “Add Video” option. AnimalTA then automatically converts the videos, allowing any format to be used without requiring external conversion tools. This process is now handled by the ffmpeg package (replacing OpenCV used in earlier versions), providing faster and more robust conversion with reduced sensitivity to corrupted frames. In addition, a multiprocessing system enables multiple videos to be converted simultaneously, significantly reducing overall processing time.

Once the videos are imported into AnimalTA, users can define a scale for each video and set “arenas” (i.e., regions of interest) within which animals are expected to appear and move. These parameters will later be used by the program during data analysis. Finally, trajectory data from external sources can be imported via the “Import Data” panel.

To facilitate data import, the current version of AnimalTA already supports data formats from several widely used video-tracking programs, including SLEAP (Pereira et al., 2022), DeepLabCut (Nath et al., 2019), and ToxTrac (Rodriguez et al., 2018). For coordinate data from these programs, users simply need to specify the file location of the tracking data to be automatically imported into the program. Data from other sources can also be imported, as long as they are stored as a single “.csv” file. In such cases, users must describe the structure of the coordinate data file through an adapted and user-friendly interface (see Figure 1).

**Figure 1.**
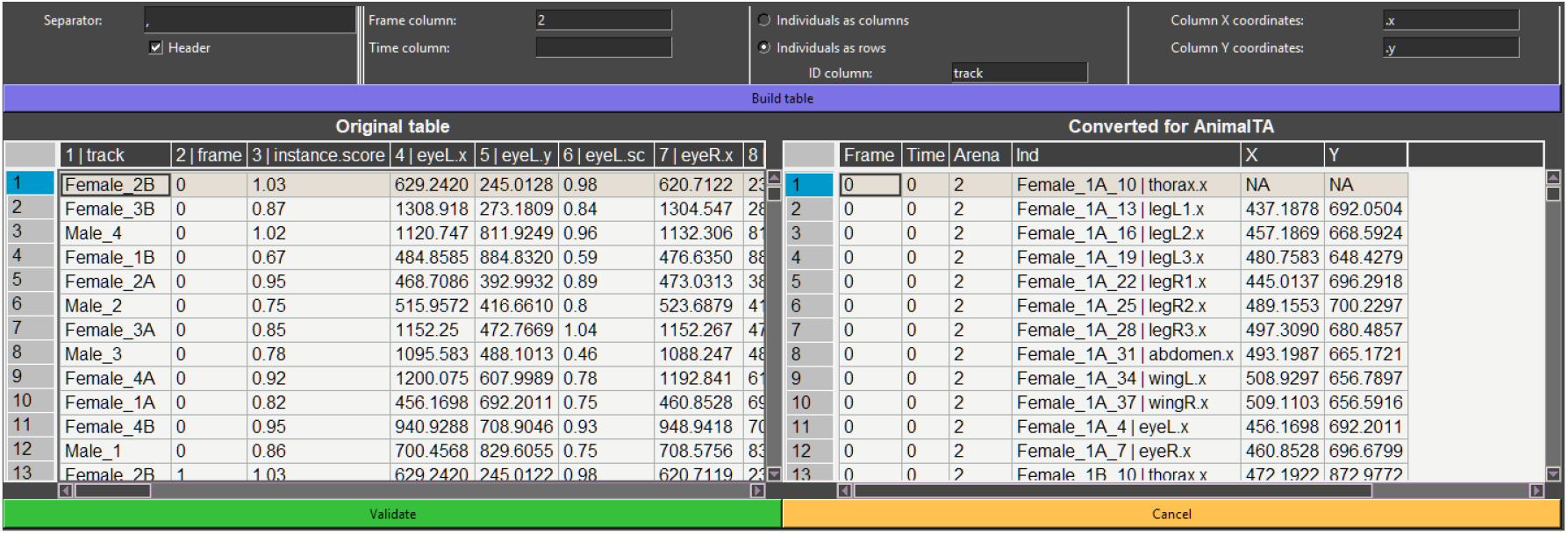
AnimalTA’s data import panel. In this panel, users can specify the structure of their original dataset using the interface at the top. The left table displays the original data, while the right table shows the converted output after configuration. Once validated, the imported data are directly integrated into the program and can be visualized, corrected, and analyzed in the same way as native AnimalTA data.

Once the data structure is defined for the first video, it is automatically saved and reused by the program for subsequent videos, so users do not need to repeat this step as long as the external trajectory data follow the same format. AnimalTA relies on the pandas and numpy packages to load, process, and save coordinate data. These final coordinate data files are stored as semicolon-delimited “.csv” files, ensuring a widely accessible and standardized format.

### 3. Improved tracking performances

In addition, AnimalTA tracking performances have been improved with the modernization of the tracking module, which now relies on a multiprocessing system (multiprocessing package, see Figure 2A for details). This update significantly enhances tracking speed, particularly for high-resolution videos or computationally intensive image processing tasks (Figure 2B–F).

**Figure 2.**
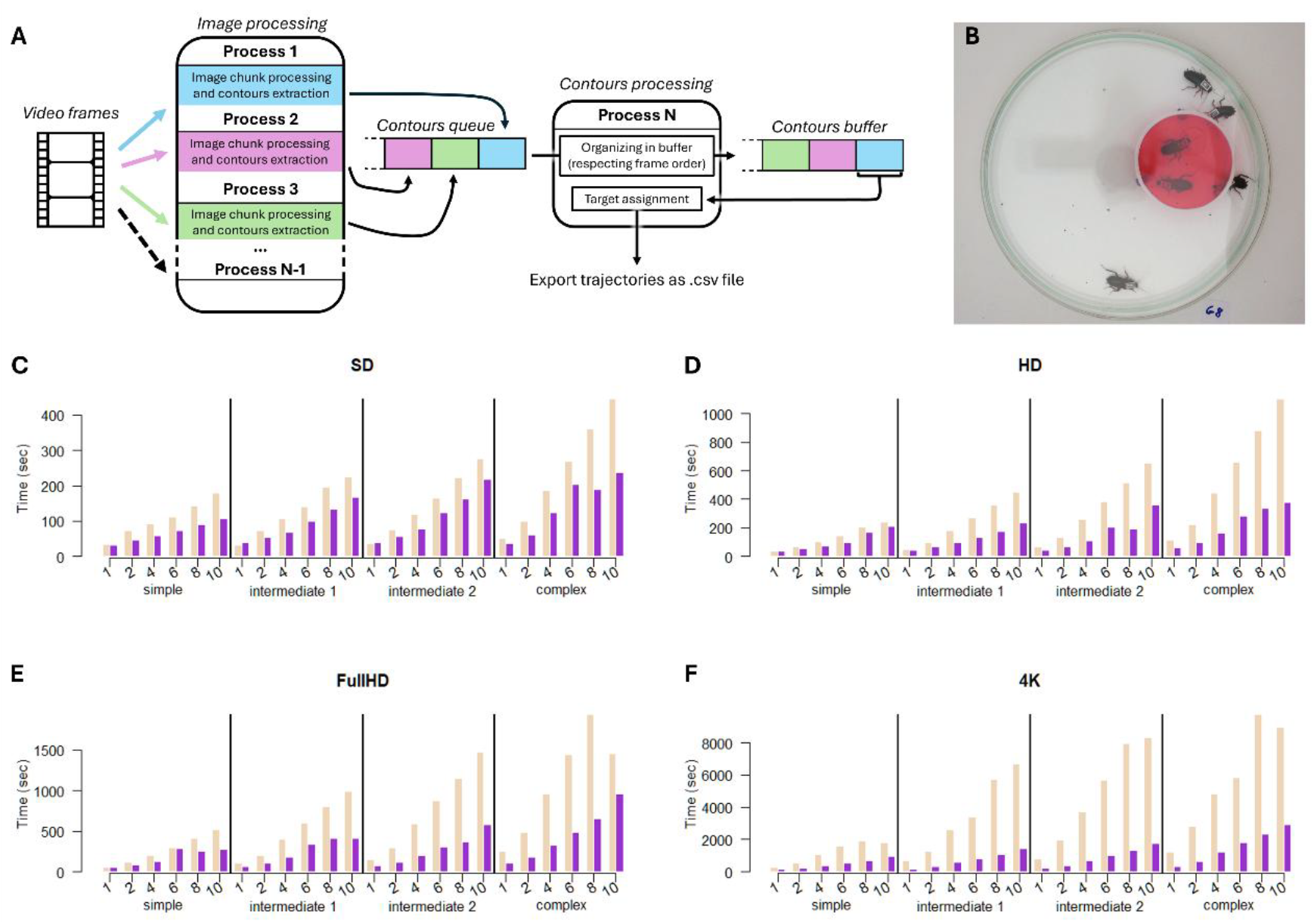
**A)** Workflow of the tracking process in AnimalTA. The tracking pipeline is divided into two main stages: image processing and contour processing. The image processing stage is the most computationally intensive part of the program. It includes extracting frames from the video, applying preprocessing steps (e.g., adjustments), and performing segmentation. Because each frame (or chunk of frames) can be processed independently, this stage can be executed in parallel and in any order. The output of this step is a set of contours (OpenCV objects), which represent the external boundaries of detected blobs (i.e., potential targets). The contour processing stage is responsible for assigning identities to these detected blobs and tracking them over time. Although this step is less computationally demanding, it must be performed sequentially and in the correct frame order, since identifying a target at frame i depends on its position at frame i–1. To optimize performance, the program uses multiprocessing. Multiple processes are dedicated to image processing, each handling different chunks of frames and pushing their results into a shared contours queue, where the frames’ data are not ordered. A separate process is dedicated to contour processing. This contour-processing process continuously monitors the queue. To reorder the frames, when specific frame data becomes available in the queue, they are retrieved and checked to determine whether they belong to the next missing frame data. If it matches, the data are processed immediately; otherwise, they are temporarily stored in an ordered buffer until the missing intermediate data arrives. Once the expected data are available, the process resumes sequential execution, emptying the buffer as much as possible before returning to the queue. This design allows the program to maximize parallel efficiency during heavy computations while preserving strict temporal order, which is required for accurate tracking. **B)** Video used to compare the efficiency of old and new versions of AnimalTA, which shows 6 female cockroaches (*Blatta lateralis*). **C to F)** Time required by the old version (light pink) and new version (purple) of AnimalTA to track the video (shown in panel B) using a computer with the following specificities: 4 cores, 8 logical processors, CPU: AMD Ryzen 5 7520U, CPU GHz: 2.8. We used the same video with different levels of resolution (4K=3840x2160px, Full HD=1920x1080px, HD=1280x720px, SD=720x480px), duration (1, 2, 4, 6, 8, and 10 min) and image processing complexity (simple=greyscale background subtraction; intermediate 1=colored background subtraction; intermediate 2=colored background subtraction and light correction; complex=colored background subtraction, light correction and stabilization). The new version is either faster or as fast as the previous version in speed, and its performance advantage grows with higher video resolution and greater task complexity. A similar comparison was repeated using a computer with different characteristics and resulted in similar findings (see Supplementary material S1).

### 4. Visualization and correction of trajectories

While the video tracking programs are becoming increasingly powerful and efficient, they are not yet perfect, and tracking inaccuracies remain common (Figure 3). In the new version of AnimalTA, special care has been taken to make data correction as fast and intuitive as possible.

**Figure 3.**
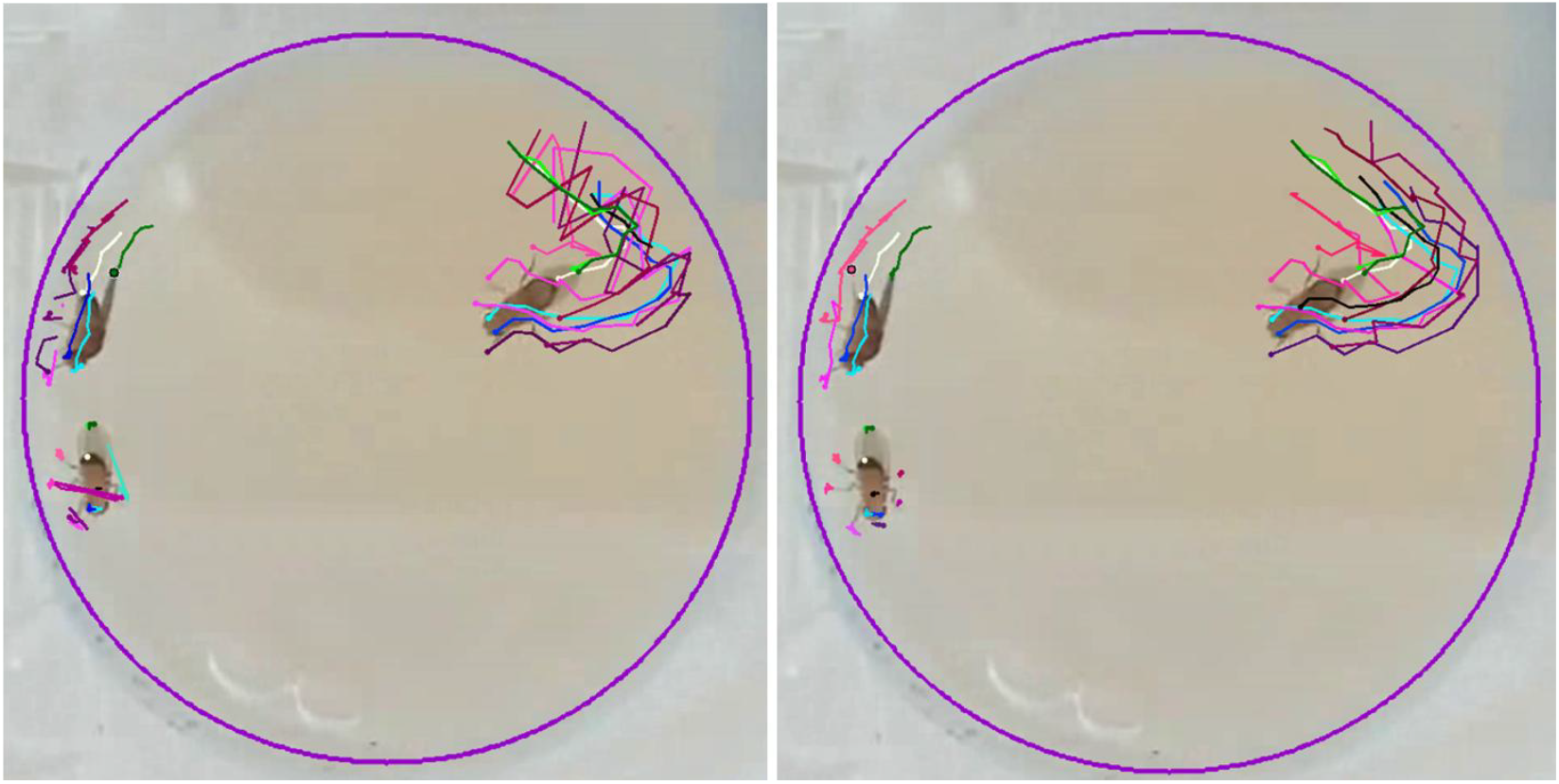
Illustration of AI-tracked groups of 3 Drosophila (left panel) and the same dataset after manual correction using AnimalTA (right panel). Tracking points were the 6 leg and 2 wing extremities, the two eyes, the thorax, and the posterior part of the abdomen for each individual. The image was provided by Aijuan Liao and Ebi Antony George, and obtained from experiments conducted in the Kawecki group at the Department of Ecology and Evolution, University of Lausanne.

AnimalTA allows for a rapid and efficient visualization of tracking data by superimposing tracked coordinates onto the original video (see Figure 3). Users can play the video at variable speeds, navigate directly to specific frames or timestamps, and zoom in or out as needed. The new version also introduces a “Lock View” option, which allows users to zoom in while the display automatically follows the target’s movement. This feature enhances the observation of fine details in moving individuals without requiring constant manual adjustment of the zoom window. To facilitate the detection of tracking errors, two additional features have been implemented: one allows users to jump directly to frames with missing coordinates, and the other highlights suspicious trajectories that exhibit abnormally high speeds relative to an individual’s typical movement patterns.

The program offers numerous tools to facilitate data correction, including the ability to fix isolated errors, interpolate across sequences of frames, perform rapid frame-by-frame corrections, and resolve identity swaps. The new version further extends these capabilities by allowing users to merge trajectories from different targets, batch-delete individuals or coordinates, and remove all coordinates within a defined spatial region of the video. It also introduces new options for easier navigation within the coordinate data frame. These new features enable researchers to flexibly adapt their use of AnimalTA to specific experimental designs and the objectives of behavioural tests.

### 5. Extension of the analysis toolset

With this update, the program introduces new analysis methods, enhancing AnimalTA’s versatility in data processing and result visualization. For example, it now allows the creation of combined heatmaps by superimposing trajectories from different arenas, as well as heatmaps based on segmented images extracted from the video. These features produce clear, informative, and visually appealing outputs (Figure 4).

**Figure 4.**
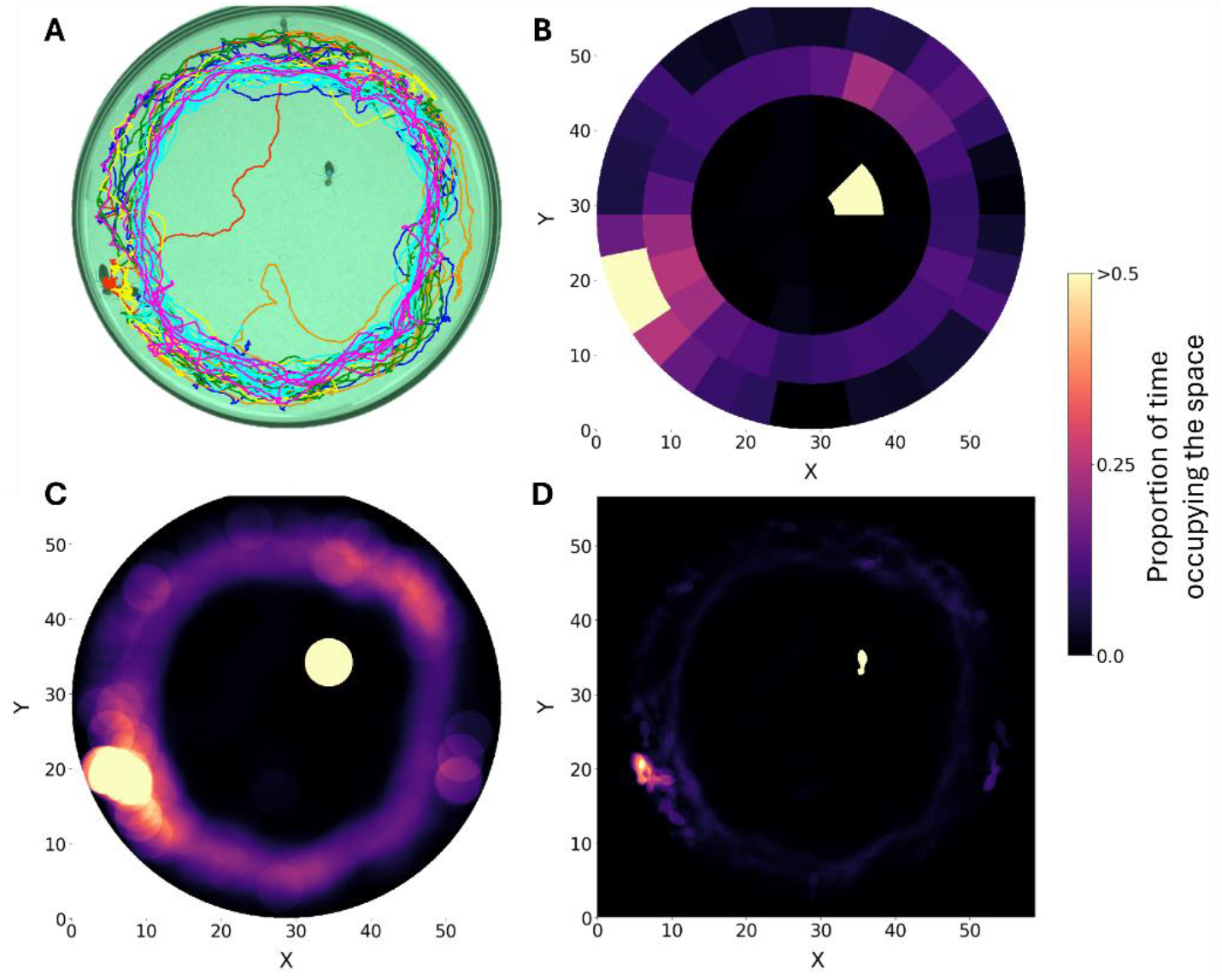
Different representations of the movement of 10 grouped termites in a 47-second video. **A)** Frame extracted from the video with all termite trajectories overlaid. **B)** Combined exploration grid derived from the same trajectories. The arena is divided into equal-sized cells, with cell color indicating the proportion of time spent by individuals within each cell (time spent by all individuals in a cell / [total video duration × number of individuals]). **C)** Individual-centered method in which termites are considered to be exploring in a circular area centered on their body position across frames. **D)** Heatmap generated from segmented video frames rather than trajectory data. All representations are exported directly from AnimalTA.

The analysis options have also been expanded with new modules, including the ability to split a video into distinct time segments, referred to as “sequences.” This feature enables users to obtain separate results for different time periods, such as the initial minutes of a recording versus the remainder, or to compare data collected before and after an individual enters a specific arena, among a wide range of possible applications. Another new option has also been introduced to calculate the duration of stops and movements performed by a target during the video. The program can either compute average values of the data or return a list of all stop and movement events observed. Such data are commonly used in survival analysis or for modelling animal locomotory behavior (Jeanson et al., 2005; Chiara et al., 2022). The last new feature is the ability to generate personalized, detailed data files containing frame-by-frame variables selected by the user. The parameters that can be extracted for each frame include distance traveled, speed, movement state (stopped or moving), corrected movement state, acceleration, orientation, change in orientation angle, angular speed, meander, distance to elements of interest, distance to other individuals, and position within the exploration grid (see Figure 4B).

## Conclusion

Since its first launch in 2023, AnimalTA has rapidly gained an increasing number of users, most of whom are animal scientists who frequently and repeatedly use this free tool to track and analyse animal movement and behaviour. Thanks to their enthusiastic support and feedback, we have been able to make an important update to AnimalTA, which greatly enhances the tool’s utility for many more users. Many existing general-purpose tracking tools and machine-learning-based tools have their own strengths in tracking ability, enabling researchers to adaptively choose the most efficient tool for their research topic, species, and experimental design. Now, with the updated AnimalTA, which allows importing and processing tracking outputs from other tracking tools, their users can also benefit from the most distinctive post-tracking functions of AnimalTA, such as rapid correction of tracking errors, versatile data analysis, and effective result visualisation.

## Acknowledgments

We are grateful to Iago Sanmartín Villar, Tristan Robineau, Srikrishna Narasimhan, Emma Chereskin, and Martin Dessart for their help with testing AnimalTA and translating program information. We also thank Ebi Antony George, Aijuan Liao, and the Kawecki group at the Department of Ecology and Evolution, University of Lausanne, for sharing their videos and tracking data on Drosophila. Finally, we thank all the people, too numerous to be cited individually, who used AnimalTA and took the time to provide feedback, which helped us improve the program. This work was supported by the research grant SONATA: 2022/47/D/NZ8/01758 funded by the National Science Center of Poland (Narodowe Centrum Nauki).

