## Supplementary Figure 1 for "AnimalTA: A simple yet flexible tool for video tracking and manual corrections"

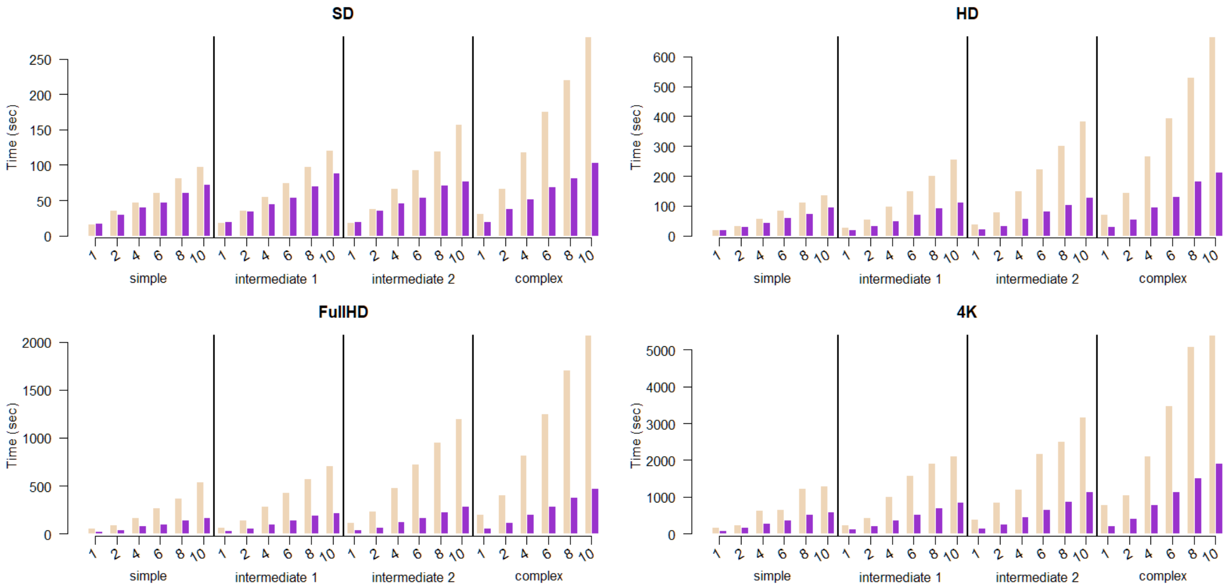


**Figure S1.** Time required by the old version (light pink) and new version (purple) of AnimalTA to track the video (shown in Figure 2, panel B) using a computer with the following specificities: 8 cores, 16 logical processors, CPU: 11th Gen Intel i7 - 11800H, CPU GHz: 2.3. We used the same video with different levels of resolution (4K=3840x2160px, Full HD=1920x1080px, HD=1280x720px, SD=720x480px), duration (1, 2, 4, 6, 8, and 10 min) and image processing complexity (simple=greyscale background subtraction; intermediate 1=colored background subtraction; intermediate 2=colored background subtraction and light correction; complex=colored background subtraction, light correction and stabilization). The new version is either faster or as fast as the previous version in speed, and its performance advantage grows with higher video resolution and greater task complexity.
